# Phosphorylation of the inner core oligosaccharide of lipopolysaccharide mediates recognition and phagocytosis of Gram-negative bacteria by Brain Angiogenesis Inhibitor 1 (BAI1)

**DOI:** 10.1101/2023.01.30.525907

**Authors:** Emily A. Billings, Linda Columbus, James E. Casanova

**Affiliations:** Department of Microbiology, Immunology and Cancer Biology, University of Virginia Health System, Charlottesville, VA 22908; Department of Chemistry, University of Virginia Health System, Charlottesville, VA 22908; Department of Cell Biology, University of Virginia Health System, Charlottesville, VA 22908

## Abstract

Recognition of lipopolysaccharide (LPS) is critical for host defense against Gram-negative bacteria and the regulation of local inflammatory responses at the host-microbial interface. We have previously shown that the adhesion G-protein-coupled receptor (aGPCR) BAI1 acts as a phagocytic pattern recognition receptor (PRR) that selectively recognizes Gram-negative bacteria through a series of five Type I thrombospondin repeats (TSRs) in its extracellular domain. Unlike Toll-like receptor 4 (TLR4), which recognizes the hydrophobic lipid A region of LPS, the BAI1 TSRs bind the core oligosaccharide of canonical LPS structures (1). In this study, we report that BAI1 requires the phosphorylated inner core L-glycero-d-manno-heptose moiety of LPS for recognition, in the context of intact, live bacteria. Mutant strains of *Salmonella enterica* serovar Typhimurium or *E. coli* BW25113 K-12 that fail to phosphorylate the inner core oligosaccharide, specifically heptose I, are not recognized by BAI1. Phosphorylation of inner core oligosaccharides is critical for membrane stability and function and is conserved across most Gram-negative species, thus allowing BAI1 to recognize a broad range of organisms.

## Introduction

Pattern recognition receptors (PRRs) have evolved to detect conserved molecular features found on and in microbes to distinguish self from non-self. These molecular motifs, defined as microbe-associated molecular patterns (MAMPs), are highly conserved across individual classes of microbes and are typically distinct from host structures. For example, exposed components of the microbial cell wall, including β-glycans and zymosan in fungi (2–4), peptidoglycan and lipotechoic acid in Gram-positive bacteria (5, 6), and LPS in Gram-negative bacteria (7, 8) are selectively recognized by distinct PRRs. The recognition of microbes in the extracellular space by humoral or cell associated receptors is critical for neutralization, initiation of inflammatory responses, and phagocytic clearance. The importance of this interaction between host and microbe is further emphasized by the targeted modification of MAMPs by pathogens and commensals for the purpose of evading and/or modulating the innate immune system.

LPS, or endotoxin, is a vital component of the Gram-negative bacterial outer membrane that contributes to membrane stability, expression and folding of outer membrane proteins, vesicle shedding, and pathogenicity (9, 10). The canonical LPS structure is composed of three regions: a highly conserved lipid A region, a core oligosaccharide, and a highly variable terminal oligosaccharide referred to as O-antigen (11–14) (Fig. 1A). LPS is synthesized sequentially, starting with the lipid A region first, then covalently linking additional sugar units to build the core oligosaccharide and finally the O-antigen regions (8, 9). Bacteria expressing the O-polysaccharide are phenotypically defined as expressing smooth LPS, while mutants lacking this structural feature are defined as rough, based on early observations of colony morphology (15). Functionally, lipid A provides an anchor within the outer membrane and confers the majority of the immunostimulatory properties of endotoxin through an interaction with TLR4 and the associated protein MD2 (8, 12, 16–18). Crystallographic studies have shown that five lipid chains of lipid A are bound by MD2 within a large hydrophobic pocket, allowing for the remaining acyl chain and the two phosphate groups from the lipid A backbone to be displayed on the surface of MD2 for presentation to TLR4 (11, 19). The phosphates play a critical role in forming hydrogen bonds and electrostatic interactions with MD2 and TLR4. In contrast, the core oligosaccharide makes minimal contact with the TLR4-MD2 complex, forming weak ionic and hydrogen bonds (19). Pathogens and gut resident microbes that modify the phosphate moieties and net negative charge of the lipid A moiety poorly stimulate TLR4 driven inflammatory responses (8, 20, 21).

**Figure 1.**
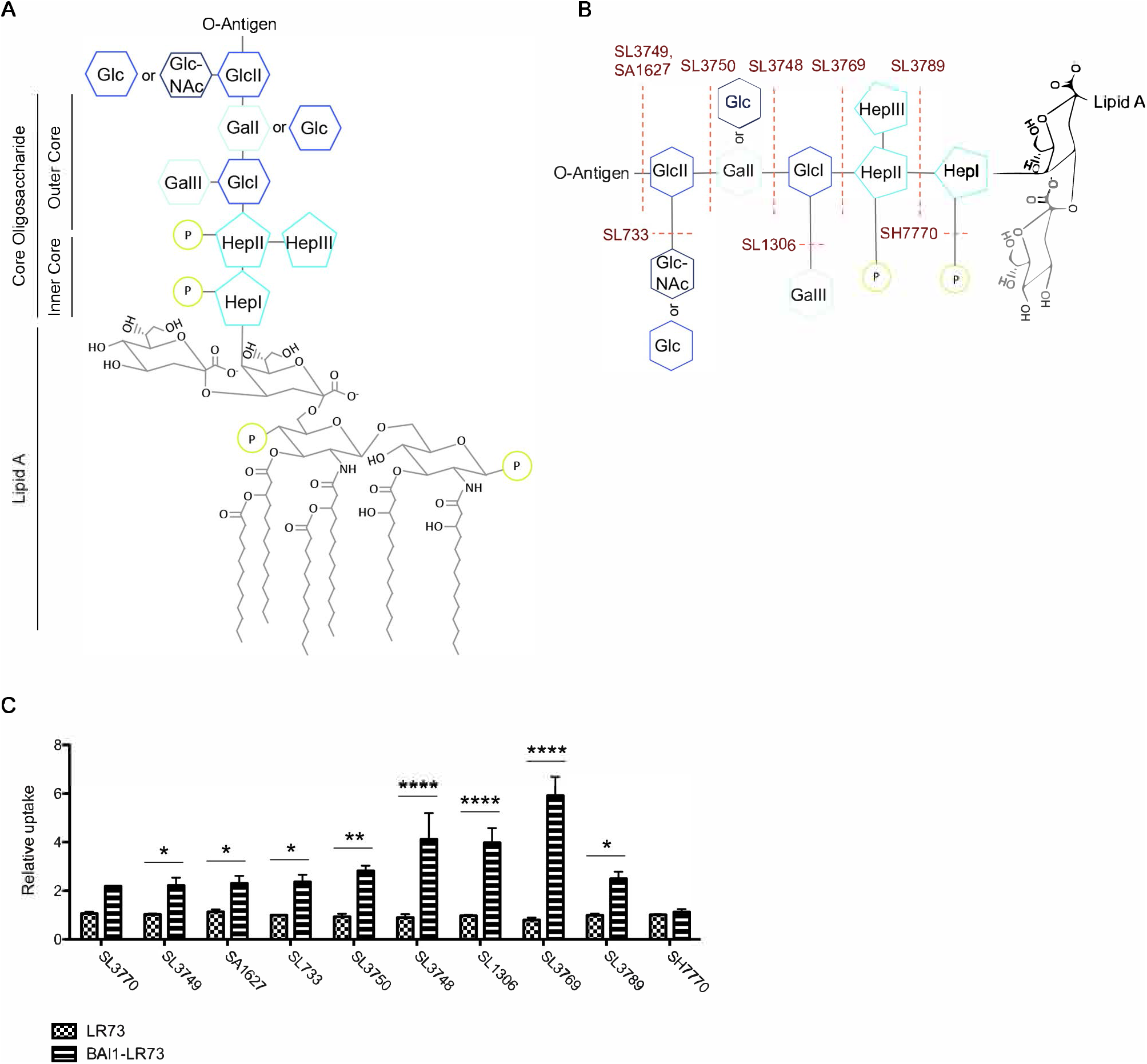
BAI1-mediated internalization of Salmonella requires phosphorylated heptose molecules in the inner core oligosaccharide. (**A**) Chemical structure of canonical enteric LPS of *S*. Typhimurium and *E. coli*. (**B**) Schematic of the core oligosaccharide structure of respective LPS mutants described in Table 1. (**C**) The internalization of the *Salmonella* strains shown in (B) was measured in parental LR73 CHO cells and cells stably expressing exogenous human BAI1 using the gentamicin protection assay as described in Materials and Methods. Data shown include the mean fold internalization ± SEM relative to control cells. The data were analyzed with 2-way ANOVA with Bonferroni post-hoc comparison. \**P* < 0.05, \*\**P* < 0.01, \*\*\**P* < 0.001. SL3770, *N* = 2; SL3749, *N* = 5; SA1627, *N* =4; SL733, *N* = 4; SL3750, *N* = 3; SL1306, *N* = 3; SL3748, *N* = 2; SL3769, *N* = 3; SL3789, *N* = 3; SH7770

The inner core oligosaccharide is also highly conserved, indicating its critical role in structural integrity. Accordingly, until very recently the innermost part of the core disaccharide backbone of lipid A, consisting of two 3-deoxy-Dmanno-otulosonic acid (KDO) moieties, was considered the minimal structure necessary for bacterial viability (8). Deep rough mutants that lack the majority of the core oligosaccharide display altered membrane composition and susceptibility to cationic peptides, detergents, and bacterial killing (22, 23). The inner core oligosaccharide consists of two KDO molecules attached to three heptose residues (Fig. 1A) (8, 24). The heptosyl backbone is covalently attached to the outer core, a trisaccharide backbone that displays more variability than the inner core (24). The highly variable O-antigen, composed of linked oligosaccharides that vary in length and composition, is attached to the outer core oligosaccharide. This region serves as a critical virulence factor, as it has been shown to inhibit phagocytosis and to promote escape from humoral and adaptive immunity for some species (25, 26).

Apart from TLR4-MD2, several other PRRs have been shown to bind LPS, with diverse consequences on host responses. For example, Caspase-11, a cytosolic component of the non-canonical inflammasome, binds the lipid A region of LPS found in the cytoplasm of infected cells to induce inflammasome activity (27–29).

Other host proteins have been shown to bind the core oligosaccharide. Surfactant protein D (SP-D) is a secreted soluble C-type lectin receptor (CLR) that recognizes the heptose moieties within the core oligosaccharide of LPS (30). Another CLR, dendritic cell-specific intercellular adhesion molecule-3 (ICAM-3)-grabbing non-integrin (DC-SIGN), recognizes the outer core oligosaccharide of LPS (31).

Thrombospondin repeats (TSRs) are conserved domains consisting of approximately 60 amino acids, found in several mammalian proteins. In other contexts, these domains are implicated in cell-cell communication and interactions with the extracellular matrix (32–34). However, several studies have found a role for TSR domains in bacterial recognition. For example, thrombospondin-1 (TSP-1), which contains 3 TSRs, has been shown to bind the peptidoglycan of several Gram-positive bacterial species resulting in adherence to host cells (35). The related protein mindin (spondin2) binds both Gram-positive and Gram-negative microbes via recognition of carbohydrate moieties in lipotechoic acid or LPS through a direct interaction with a single type I TSR (36, 37). Common features found in many of these mammalian proteins that mediate microbial recognition include positive charge, hydrophobicity, and amphipathicity (21).

BAI1 is a member of subgroup VII of the adhesion-type GPCRs that acts as a phagocytic receptor for both apoptotic cells and bacteria (1, 38–40). A key structural feature of BAI1 is the presence of five type I TSRs in its extracellular domain. Consistent with the ability of TSR domains in other mammalian proteins to bind bacterial membrane components, work from our laboratory showed that the BAI1-TSRs recognize LPS (1).

However, unlike mindin (spondin2) and TSP-1, BAI1 does not recognize Gram-positive bacteria or peptidoglycan (1). Recognition of LPS is independent of O-antigen, but dependent on the core oligosaccharide (1). However, the importance of this interaction to the innate immune response is not known. Moreover, the specific features of LPS that are recognized by BAI1 remain unidentified.

In this study, we determined that the inner core oligosaccharide is critical for BAI1-mediated recognition. Consistent with our previous findings, BAI1 was able to recognize bacteria expressing Ra-type LPS containing the inner core oligosaccharides. Importantly, BAI1-mediated recognition of bacteria required phosphorylation of the inner core heptoses, suggesting that electrostatic interactions are an important aspect of the recognition mechanism. The specificity of BAI1 for the phospho-heptose sugars in the inner core oligosaccharide suggests broad importance for recognition of Gram-negative bacteria, as this region is conserved across many commensal and pathogenic bacteria alike.

## Results

### The outer core oligosaccharide of Salmonella is dispensable for BAI1 recognition

We previously determined that BAI1 mediates the binding and internalization of non-invasive *Salmonella enterica* serovar Typhimurium, a representative Gram-negative bacterial pathogen, in several cell culture model systems (1). In a solid-phase binding assay using purified components, a soluble fragment of the BAI1 ectodomain containing the five TSRs bound to both smooth LPS (containing the polysaccharide O-antigen) and rough (Ra chemotype) LPS, lacking O-antigen. However, LPS isolated from a deep rough mutant (Re chemotype) was not recognized, indicating that BAI1 selectively recognizes one or more components of the core oligosaccharide. Similarly, in a cell-based assay we showed that fibroblasts (LR73 CHO cells) expressing exogenous BAI1 efficiently internalized *E. coli* strain DH5α, which expresses rough (Ra chemotype) LPS (39). However, the specific structural features of the core oligosaccharide that are recognized by BAI1 have not yet been determined.

To further examine the specificity of this interaction, we made use of a set of *S. typhimurium* strain LT2 mutants that lack either O-antigen or individual components of the core oligosaccharide. An enzyme required for a specific step in synthesis of the LPS core is deleted in each strain. Since synthesis of LPS occurs sequentially, these mutations can be ordered in a series resulting in a progressively truncated LPS structure.

Table 1 lists the strains used in this study and their corresponding LPS chemotypes. A diagram indicating the site at which each mutant affects core oligosaccharide structure is provided in Fig. 1B. For this analysis we used fibroblasts stably expressing exogenous human BAI1, with non-expressing parental cells as controls (Fig. S1). We also used live bacteria, to ensure that recognition by BAI1 occurs in the context of an intact bacterial membrane. Cells were incubated with bacteria at an MOI of 50 at 37°C for 30 min to allow binding and internalization. Internalization was then quantified using a standard gentamicin protection assay. Consistent with our previous observations, cells expressing BAI1 internalized more wild type *S. enterica* serovar Typhimurium (strain SL3770, which contains O-antigen) than control cells that did not express BAI1 (Fig. 1C). Internalization of two rough strains lacking O-antigen (SL3749, SA1627) was quantitatively similar to the WT, smooth strain. Similarly, LPS mutants expressing truncations in the outer core oligosaccharide were also efficiently internalized by BAI1-expressing cells. This included strains lacking N-acetyl glucosamine side chains (SL733) or the terminal glucose moiety GlcII (SL3750). Interestingly, strains expressing LPS truncated before the second hexose moiety in the outer core (SL3748), the proximal glucose moiety GlcI (SL3769) or its attached galactose, GalII (SL1306) exhibited a relative increase in BAI1-dependent internalization efficiency, suggesting that the outer core may partially mask a structural component required for binding to the TSRs (Fig. 1C). A similar increase in binding was previously reported for the mannose binding lectin (MBL) upon removal of outer core oligosaccharides (41). Collectively, these results indicate that the outer core oligosaccharide of *S. enterica* serovar Typhimurium is not significantly recognized by BAI1.

### BAI1-mediated recognition of S. typhimurium requires phosphorylated heptoses in the inner core oligosaccharide

We next analyzed the contribution of inner core oligosaccharide mutants to BAI1-mediated recognition. In contrast to the outer core sugars, which are hexoses, the inner core sugars are heptoses, two of which (HepI and HepII) are phosphorylated. As shown in Fig. 1C, loss of the two distal heptoses (HepII and HepIII, strain SL3789) resulted in decreased internalization efficiency relative to complete loss of the outer core (SL3769). However, internalization efficiency remained quantitatively similar to strains expressing the more distal outer core sugars. In contrast, loss of the most membrane-proximal heptose (HepI, strain SL1102) or its covalently attached phosphate (SH7770) resulted in complete loss of recognition by BAI1 (Fig. 1C).

Collectively, these data suggest that the distal inner core heptoses (HepII and HepIII) contribute to recognition by BAI1, but that phosphorylation of the membrane proximal heptose (HepI) is essential.

### Specificity of the BAI1-TSR domains for phosphorylated inner core heptose sugars is recapitulated in an isogenic E. coli K-12 mutant collection

Although the *Salmonella* LPS mutants utilized in Figure 1 are all derived from the parental strain *S. Typhimurium* LT2, they are not isogenic. This could result in altered behavior and interactions with host cells, independent of LPS structure. To expand upon the observations made with *Salmonella*, we used a set of *E. coli* K-12 BW25113 LPS mutants derived from the Keio collection, a publically available single-gene deletion mutant library. Table 2 lists these strains and their corresponding LPS structures. The parent strain, K-12 BW25113, lacks O-antigen and is phenotypically rough.

As we observed for *Salmonella*, expression of BAI1 significantly enhanced recognition of the parental *E. coli* K-12 BW25113 strain, and this was further enhanced with a strain lacking the outer core oligosaccharide (rfaG) (Fig. 2, A and B). However, unlike *Salmonella*, loss of the membrane-distal heptoses HepII and HepIII (rfaF) or the phosphate attached to HepII (rfaY) reduced binding to near background levels. Similarly, removal of the phosphate on the membrane-proximal HepI (rfaP) completely blocked recognition by BAI1. Together, these observations suggest that phosphorylation of both HepI and HepII is essential for recognition of *E. coli* by BAI1.

**Figure 2.**
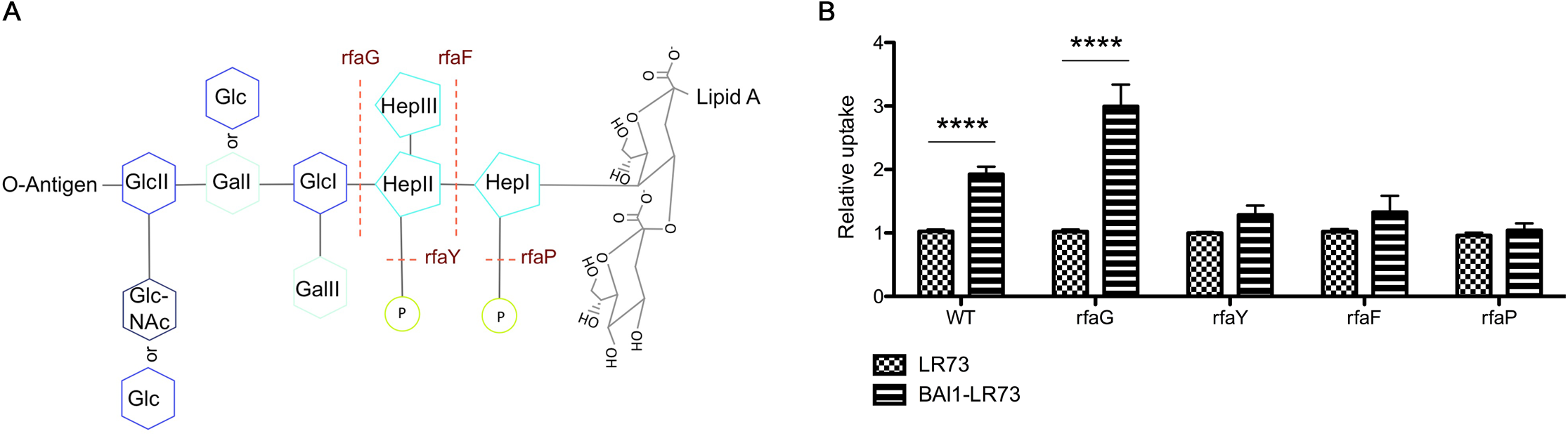
BAI1-mediated phagocytosis of E. coli K-12 requires phosphorylated heptose sugars in the inner core oligosaccharide. (**A**) Schematic of the core oligosaccharide structure with the location of respective LPS mutants described in Table 2. (**B**) Uptake of the *E. coli* LPS mutants shown in (**A**) was measured in BAI1-expressing LR73 CHO cells and control cells using the gentamicin protection assay as describe above. Data indicate the mean fold increase ± SEM normalized to control (non-BAI1-expressing) cells. Data were analyzed with 2-way ANOVA with Bonferroni post-hoc comparison. \**P* < 0.05, \*\*\**P* < 0.001, \*\*\*\**P* < 0.0001. WT, *N* = 14; ΔrfaG, *N* = 6; ΔrfaF, *N* =7; ΔrfaP, *N* = 13

### Structural and biophysical properties of the TSRs are consistent with charge interactions driving recognition of LPS

Analysis of the structure of the type I TSRs reveals biophysical properties that may contribute to the mechanism of recognition and specificity of BAI1 for the inner core oligosaccharide of LPS. The highly conserved TSR domains consist of ∼60 amino acids, which form three anti-parallel strands that interact by interlacing and stacking arginine and tryptophan side chains (42, 43). Figure 3 shows homology models of the five BAI1 TSR domains generated by threading the sequence of the BAI1 TSRs onto the known structure of TSR1 from thrombospondin-1 (pdb id 1LSL) using SWISS-MODEL (42). Calculated electrostatic maps (ref) reveal a positively charged surface on one face of the domain that likely mediates electrostatic interactions with its negatively charged ligands (37, 42, 44–46). Interestingly, the five TSRs in BAI1 have variable intensities centered at different locations with respect to the most electropositive region, suggesting that the different domains may differentially interact with ligands (Fig. 3).

**Figure 3.**
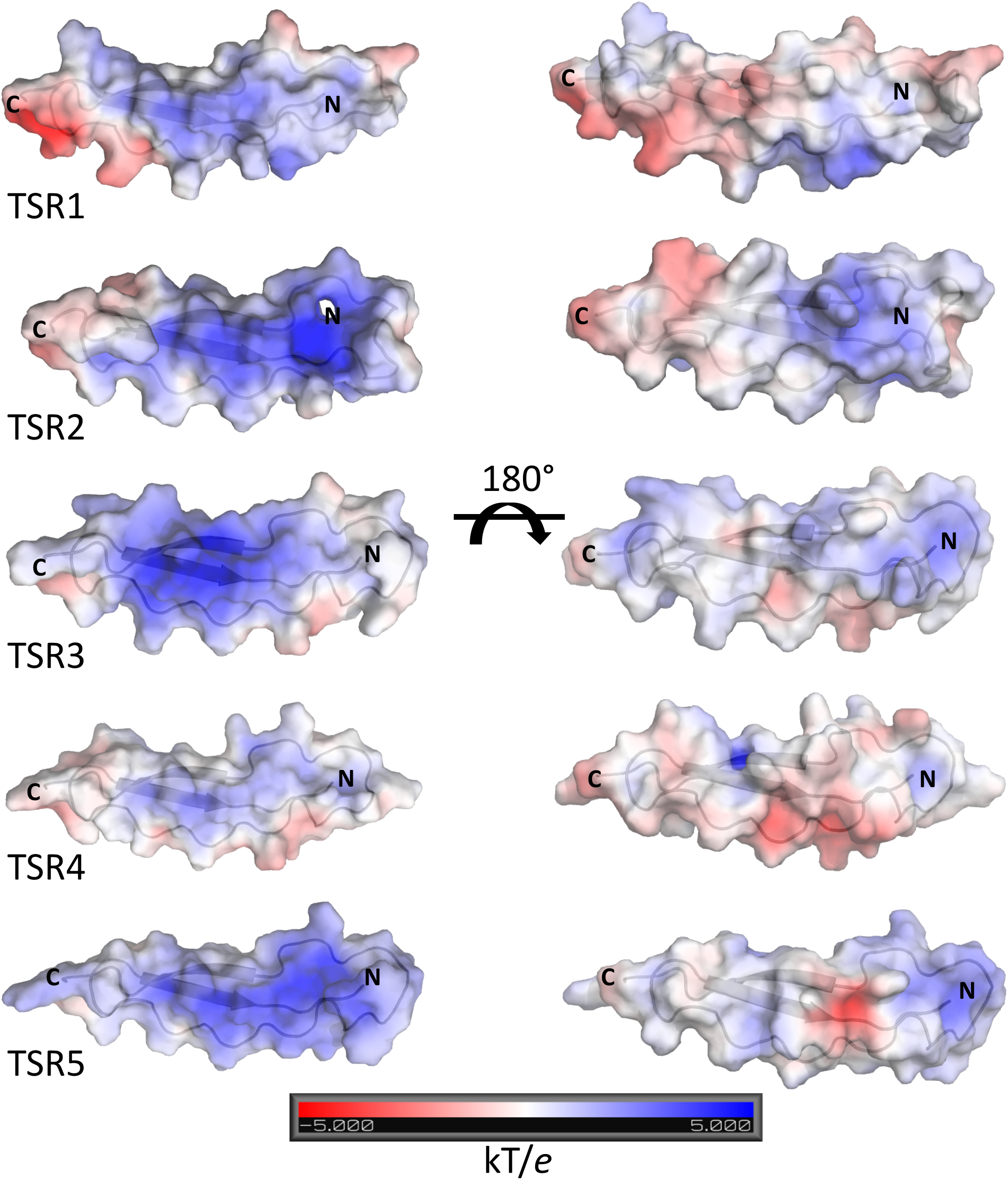
Homology models of the five BAI1 thrombospondin repeats (TSRs). For each TSR, the BAI1 sequence was modeled based on the crystal structure of the TSR1 of thrombospondin-1 (db id 1LSLS). The electrostatic surface potential is shown as a transparent surface with the protein backbone rendered as cartoon. The N- and C-termini are labeled. Note that TSRs adopt an elongated structure comprised of three interacting - strands. One surface of each TSR is rich in positively charged side chains (as evident from the electropositive surface; blue), while the opposite surface has a more electroneutral character.

## Discussion

We recently identified BAI1 as a novel PRR that directly binds and drives the internalization of Gram-negative bacteria through its recognition of LPS (1). Ligation of BAI1 by bacteria or soluble LPS also triggers activation of the macrophage NOX2 NADPH oxidase complex, thereby enhancing the killing of internalized bacteria (39).

Recognition of LPS is direct and is mediated by interaction between the five Type I TSRs in the extracellular domain of BAI1 and the core oligosaccharide region of LPS (1). Importantly, this mechanism of LPS binding is distinct from the TLR4-MD-2 complex, which binds to the acyl chains and di-glucosamine backbone of LPS (11). In this study we used mutant strains of *S. Typhimurium* and *E. coli* deficient in each of the steps of LPS oligosaccharide synthesis to map the structural determinants that are recognized by BAI1. Using live bacteria, we show that the outer core oligosaccharide is dispensable for binding to BAI1. In contrast, phosphorylation of one or more of the L-glycero-d-manno-heptose residues in the inner core oligosaccharide is essential for recognition of LPS by the BAI1 TSR repeats.

Whether phosphorylation of both HepI and HepII is required for TSR binding is still unclear and may differ between *E. coli* and *Salmonella*. In *E. coli*, phosphorylation of HepI by the kinase WaaP/RfaP is required for subsequent phosphorylation of HepII by a second kinase, WaaY/RfaY, such that WaaP/RfaP mutants lack heptose phosphorylation entirely (47, 48). However, our data indicate that an *E. coli* RfaY mutant is very poorly recognized by BAI1, suggesting that phosphorylation of HepII alone is sufficient to mediate binding to BAI1. In contrast, HepII does not appear to be essential for recognition of *Salmonella* by BAI1. Although our *Salmonella* mutant collection did not include an RfaY equivalent, a mutant completely lacking HepII and HepIII (SL3789) was still recognized by BAI1, as efficiently as the wild type LT2 strain. In this case, phosphorylation of HepI appears to be more important than HepII. The reasons for these discrepancies are not known, but could relate to the precise orientation of the heptose moieties in the bacterial membrane and the degree to which they are exposed at the cell surface.

This study provides insight into the specificity of the TSR domains for bacterial LPS, but many outstanding questions remain. The importance of the phosphorylation of the inner core oligosaccharide for viability and fitness of many Gram-negative bacteria indicates that, like TLR4, BAI1 recognizes an essential feature of LPS. As mentioned above, BAI1 also mediates the clearance of apoptotic cells by macrophages, through recognition of exposed, negatively charged phosphatidylserine by its TSRs (38). Interestingly, the cellular response to bacterial recognition is distinctly pro-inflammatory, while the response to apoptotic cells is anti-inflammatory. Whether these differences are due to co-ligation of other receptors (e.g. TLR4) or to differential coupling of BAI1 to downstream signaling machinery remains to be determined but will be an important area of future research.

## Methods and Materials

### Isolation and culture of cells

LR73 CHO cell lines have been described previously (38) and were cultured in αMEM, (Gibco) containing 10% FBS and 1% pen-strep under selection with puromycin to maintain BAI1 expression (Invitrogen).

### Bacterial strains and culture

All bacteria were cultured overnight in LB broth under aerobic conditions before use. *E. coli* BW25113 parent strain and LPS mutants listed in Table 2 and Figure 2A were from the *E. coli* Genetic Stock Center Keio collection (49). The *Salmonella enterica* serovar Typhimurium LT2 wild type and LPS mutants, originally from the *Salmonella spp*. Genetic Stock Center, were a gift from Dr. Joanna Goldberg at Emory University.

### Gentamicin protection assay

The gentamicin protection assay was performed as described previously (1). Briefly, 5 × 10^4^ LR73 cells/well were seeded into 24-well plates 18 hours before infection. Cells were incubated with bacteria at an MOI of 50 for 30 min at 37°C in αMEM containing 10% heat-inactivated FBS, after spinning bacteria onto the cells at 500 x g for 5 min at 4°C to synchronize uptake. After 30 min of internalization, cells were treated with gentamicin (500 μg/ml, Gibco) for 60 min to kill extracellular bacteria, but leave intracellular bacteria viable. Bacterial internalization was measured by colony forming assay in Hank’s balanced salt solution (HBSS) plus calcium and magnesium and 0.5% saponin.

## Figure Legends

**Table 1.** *List of wild type and LPS mutant S. Typhimurium strains used in this study.* For each strain, the LPS chemotype or phenotype is provided, along with a description of the mutation in the LPS structure.

**Table 2.** *List of wild type and mutant E. coli strains used in this study.* All strains are variants of K-12 BW25113. For each strain, the LPS chemotype or phenotype is provided along with a description of the mutation in the LPS structure.

**Figure S1.** Measurement of protein expression in LR73 control cells or cells stably expressing BAI1 LR73 control cells and cells exogenously expressing BAI1 are maintained under puromycin selection. Cell lysates were probed with an anti-BAI1 antibody (Allele Biotechnology, ABP-PAB-10421) to confirm maintenance of BAI1 expression. BAI1 is detected as two bands in LR73 cells.

## Notes

### Competing Interest Statement

The authors have declared no competing interest.

